# Sounding the alarm: sex differences in rat ultrasonic vocalizations during Pavlovian fear conditioning and extinction

**DOI:** 10.1101/2022.09.13.507801

**Authors:** MA Laine, JR Mitchell, J Rhyner, R Clark, A Kannan, J Keith, MC Pikus, E Bergeron, I. Ravaglia, E Ulgenturk, A Shinde, RM Shansky

## Abstract

Pavlovian fear conditioning is a prevalent tool in the study of aversive learning, which is a key component of stress-related psychiatric disorders. Adult rats can exhibit various threat-related behaviors, including freezing, motor responses and ultrasonic vocalizations (USVs). While these responses can all signal aversion, we know little about how they relate to one another. Here we characterize USVs emitted by male and female rats during cued fear acquisition and extinction and assess the relationship between different threat-related behaviors. To probe the effects of aversive stimulus intensity, we exposed the rats to mild (0.3 mA), moderate (0.5 mA) or strong (1 mA) foot shocks. We found that males consistently emitted more alarm calls than females, and male alarm calls were more closely contingent on shock intensity than were female alarm calls. Furthermore, 25 % of males and 45 % of females did not emit alarm calls. Males that made alarm calls had significantly higher levels of freezing than males who did not, while no differences in freezing were observed between female alarm callers and non-callers. Alarm call emission was also affected by the predictability of the shock; when unpaired from a tone cue, both males and females started emitting alarm calls significantly later. Some rats continued to alarm-call during extinction learning (90% of males, 30% of females) and retrieval (65% of males, 20% of females). Collectively these data suggest sex-dependence in how behavioral readouts relate to innate and conditioned threat responses. Importantly, we suggest that the same behaviors can signal sex-dependent features of aversion.

**Significance statement:** Behavioral neuroscientists can access various outputs during behavioral tests to draw conclusions about internal states of animals. While freezing is the most common index of rodents feeling threatened, these animals also emit specific ultrasonic vocalizations during aversive situations. Here we record several motor and vocal behaviors to assess how they relate to each other as threat responses, and how such relationships vary across sex. We found robust differences in how much male and female rats engaged in so-called alarm vocalizations. These vocalizations were subject to extinction in both sexes, but correlated with freezing only in males. As the field advances to include more females in preclinical research, it is crucial that we understand how similar-appearing outputs may reflect sex-biased features.

## Introduction

Understanding how memories of aversive situations are formed is an important goal for preclinical research on post-traumatic stress disorder (PTSD) and other diagnostic categories where such memories are affected (de Quervain et al., 2009; Giustino & Maren, 2015; Heldt et al., 2007). These processes have been significantly elucidated by preclinical approaches using animal models. Most notably, Pavlovian fear conditioning has long been the gold standard method for studying the acquisition and extinction of associations between aversive unconditioned stimuli (US, typically a mild electric foot shock) and previously-neutral conditioned stimuli (CS, such as an auditory tone or a scent). After repeated CS-US paired presentations, mere exposure to the CS begins to elicit defensive behaviors (Bolles & Collier, 1976; Fanselow, 1984; Killcross et al., 1997; LeDoux, 2000). Further repeated exposure to the CS in the absence of the US typically results in extinction of the defensive behavior, a process which has been leveraged to improve exposure therapies for humans (Davis et al., 2006; Hofmann et al., 2006; Ressler et al., 2004). In fear conditioning experiments, the typical behavioral readout of increased association between the CS and US is locomotor behavior, such as rapid darting or the complete absence of movement, i.e. freezing (Blanchard et al., 1986; Bolles & Collier, 1976; Borkar et al., 2020; Fadok et al., 2017; Fanselow, 1980; Gruene et al., 2015; Mitchell et al., 2022).

Another facet of the rodent threat response is ultrasonic vocalization (USV). USVs have been described across the rodent lifespan as potential indicators of emotional valence (Knutson et al., 2002) and as a means of communicating among pups and dams (Granata et al., 2021; Hofer & Shair, 1978; Portfors, 2007; Wöhr & Schwarting, 2008) as well as adults (Kalenscher et al., 2021; Reinhold et al., 2019; Takahashi et al., 2010). When faced with aversive situations, such as exposure to predator odors or stressful behavioral tests (Borta et al., 2006; Brudzynski & Ociepa, 1992; Fendt et al., 2018), rats emit specific low-frequency (around 22 kHz) calls, often termed “alarm calls”. As expected, fear conditioning using foot shocks robustly produces alarm calls in both male and female rats from various genetic backgrounds (Dupin et al., 2019; Schwarting et al., 2007; Wöhr et al., 2005). Quantification of these calls has revealed some variations in both total amounts produced across experiments, and in auditory parameters (Willadsen et al., 2021; Yee et al., 2012). However, precise temporal patterns of call emission throughout fear conditioning and extinction, and their relationship to other threat-associated behaviors remain understudied. Such behavioral readouts represent potentially fruitful avenues for capturing a more multi-faceted picture of learned fear in both sexes.

Our aim was to provide a comprehensive characterization of USVs across both conditioned fear acquisition and extinction, covering the dynamic ways in which vocalizations change across these testing sessions. Additionally, we examined how unconditioned stimulus intensity (i.e. foot shock intensity) and predictability moderated behavioral readouts. The objective of this work was to expand our understanding of what USVs can tell us about affective states, and whether these patterns differ between male and female rodents. As we continue to normalize the use of female rodents in behavioral neuroscience, it is critical to know if the same experimental measures collected for decades using only males reflect the same emotional states in females, or whether behavioral repertoires are sex-biased (Bangasser & Cuarenta, 2021; Rechlin et al., 2022; Shansky & Murphy, 2021).

## Methods

### Animals

Male (average weight 445.87 g at testing, total N = 101) and female (average weight 267.32 g at testing, total N = 99) Sprague-Dawley rats were purchased from a commercial breeder (Charles River, MA), and acclimated to the vivarium for 7 days before the start of handling. The vivarium was temperature (22 °C ±1 °C) and humidity (40 % ±10 %) controlled on a 12h light-dark cycle (lights on: 7:00 am), and all rats were pair-housed with ad libitum access to food (RMH 3000, Purina) and water (filtered tap water). Each cage contained a tinted plexiglass chamber for nesting and enrichment, and heat-treated pine shavings for bedding. All behavioral experiments were conducted during the light phase between 9:00 am and 3:00 pm. All procedures were conducted in accordance with the National Institutes of Health Guide for the Care and Use of Laboratory Animals and were approved by the Northeastern University Institutional Animal Care And Use Committee.

### Behavioral data collection and analysis

Rats were handled on two days prior to cued fear conditioning, and habituated to transport from the vivarium to the testing room on a trolley the day before conditioning. On the conditioning day, rats were brought in 30 minutes before the start to habituate to the testing room (< 15 meters from the vivarium) and ambient noise. They were then placed in sound-attenuating conditioning chambers (H10-24A housing a Rat Test Cage, Couldbourn Instruments, Allentown, PA), consisting of plexiglass and metal walls and a metal grid floor for the delivery of foot shocks (H10-11R-TC Shock Floor, Coulbourn Instruments). The chambers were dimly lit by an overhead light (2 lux). Following a 5-minute baseline period with no stimulus presentations, the rats were sequentially exposed to a total of 7 CS-US pairings (Fig 1A). As CS, we used a 30 second 7 kHz tone played in each chamber by a speaker at approximately 75 dB. For paired fear conditioning (Fig 1, 2, 4–7) the foot shock US, lasting 0.5 seconds, was presented at the end of the tone, with the two stimuli co-terminating. Each rat received foot shocks representing one of three shock intensities: 0.3 mA (mild shock intensity), 0.5 mA (moderate shock intensity) or 1 mA (high shock intensity). The inter-trial interval (ITI) was of varying length between each of the CS-US pairings (90-330 seconds). Two to four rats were conditioned simultaneously in the same room, without mixing sexes within each run. Chambers and the testing cages were cleaned with water and ethanol between each run. See figure legends for exact Ns for each group.

**Figure 1.**
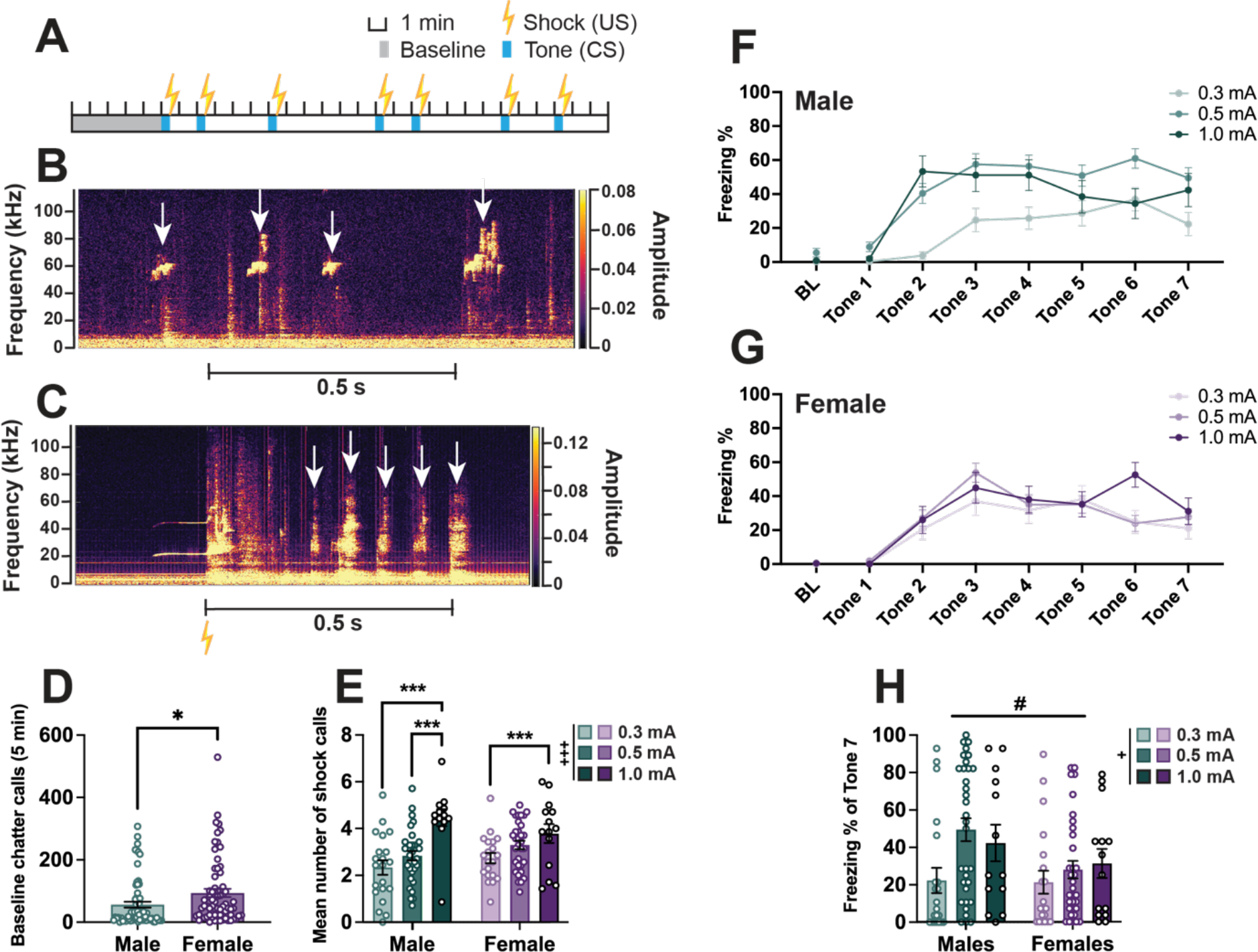
Sex differences in ultrasonic baseline and shock calls, and freezing during fear conditioning. **A**. Schematic of cued fear conditioning showing timing and duration of tones (CS) and shocks (US). **B**. Representative spectrogram (from DeepSqueak) showing typical high-frequency ultrasonic calls (white arrows) recorded during the baseline period. **C**. Representative spectrogram showing typical shock calls (white arrows). Yellow lightning symbol denotes shock onset time. **D**. Bar graph showing the total number of baseline chatter calls emitted by male and female rats prior to fear conditioning. **E**. Bar graph showing the mean number of shock calls emitted in response to each shock averaged across the trial (7 shocks) by each animal, split by sex and shock intensity. **F**. Percent of time male rats within each shock intensity group spent freezing during baseline (BL, first 2 minutes) and each tone. **G**. Percent of time female rats within each shock intensity group spent freezing during baseline (BL, first 2 minutes) and each tone. **H**. Comparison of freezing percent at the end of fear conditioning (Tone 7) between males and females, and across shock intensities. Bar graphs depict mean ± SEM, and each dot represents a single animal. Symbols along line graphs indicate mean ± SEM. Significant main effects of shock intensity (+) and sex (#), and post hoc/pairwise comparisons (*) are denoted with different symbols, with either 1, (*p* < 0.05), 2 (*p* < 0.01) or 3 (*p* < 0.001) symbols depicting degree of significance.

**Figure 2.**
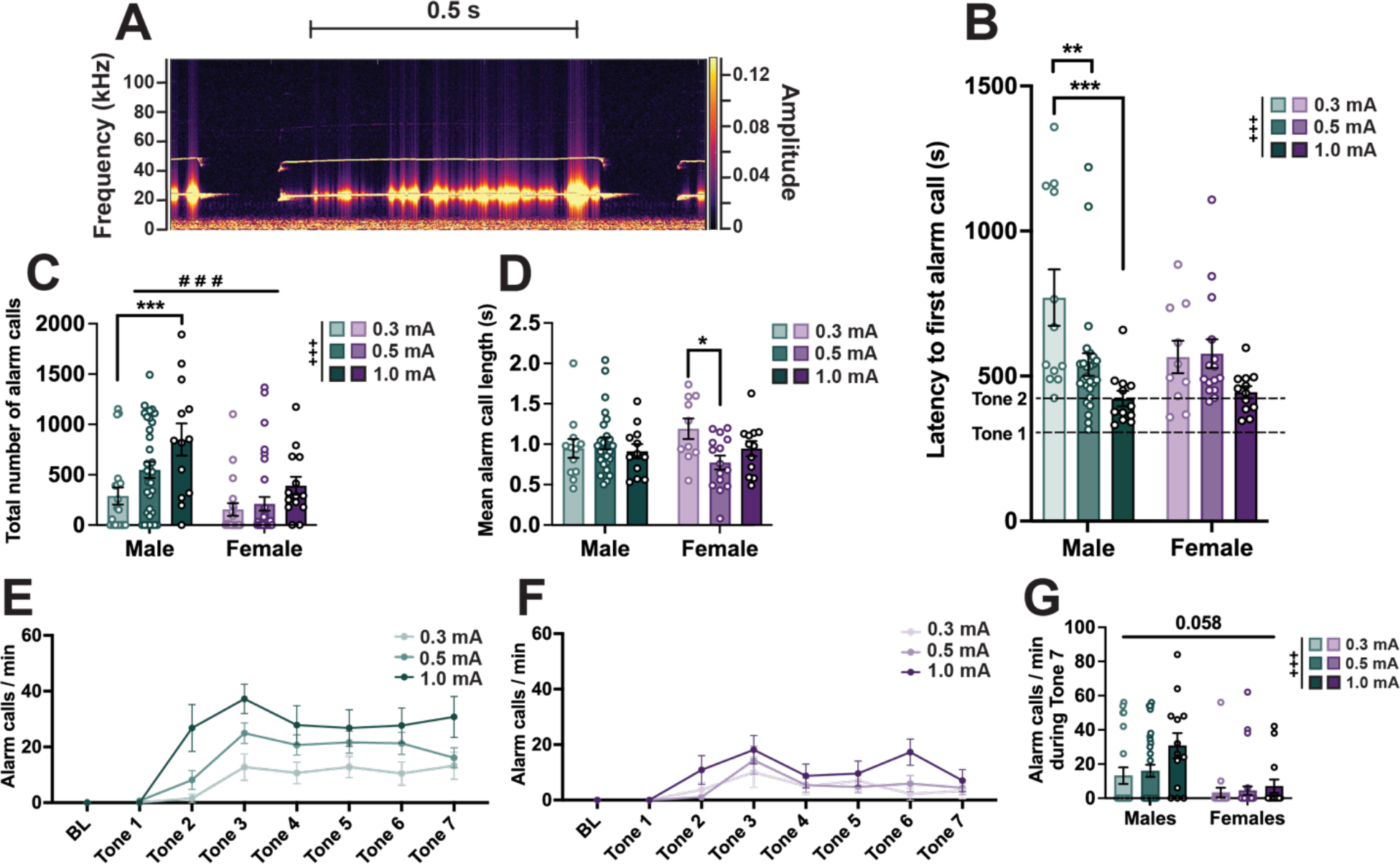
Sex differences in ultrasonic alarm calls during fear conditioning. **A**. Representative spectrogram (from DeepSqueak) showing an ultrasonic alarm call, with a principal frequency of approximately 22 kHz. **B**. Bar graph depicting the latency of each animal to emit their first alarm call, split across sex and shock intensity. Dashed lines indicate the timing of the first and second tone starts. **C**. Bar graph depicting the total number of alarm calls emitted during a fear conditioning trial, split across sex and shock intensity. **D**. Bar graph depicting the mean alarm call length, split across sex and shock intensity. **E**. Line graph showing the normalized (per minute) alarm call rate of male rats across shock intensity groups during baseline (BL, 5 minutes) and each tone. **F**. Line graph showing the normalized (per minute) alarm call rate of female rats across shock intensity groups during baseline (BL, 5 minutes) and each tone. **G**. Comparison of alarm call rate at the end of fear conditioning (Tone 7) between males and females, and across shock intensities. Ns: C, E – G: 67 males (0.3 mA = 21, 0.5 mA = 33, 1 mA = 13), 67 females (0.3 mA = 20, 0.5 mA = 33, 1 mA = 14); B & D: 50 males (0.3 mA = 12, 0.5 mA = 26, 1 mA = 12), 37 females (0.3 mA = 10, 0.5 mA = 15, 1 mA = 12) (Non-alarm callers excluded, see Fig 5 for analysis of Alarm callers vs Non-alarm callers). Bar graphs depict mean ± SEM, and each dot represents a single animal. Symbols along line graphs indicate mean ± SEM. S Significant main effects of shock intensity (+) and sex (#), and post hoc comparisons (*) are denoted with different symbols, with either 1, (*p* < 0.05), 2 (*p* < 0.01) or 3 (*p* < 0.001) symbols depicting degree of significance.

A distinct cohort of animals were part of an experiment to compare exposure to unpaired (16 males + 16 females) or paired (18 males, 16 females) fear conditioning (Fig 3). The parameters of the CS and US were identical to paired fear conditioning, aside from the shock always occurring a minimum of 60 seconds after the cessation of the tone. The ITIs of CS and US presentations were varied (Fig 3A).

**Figure 3.**
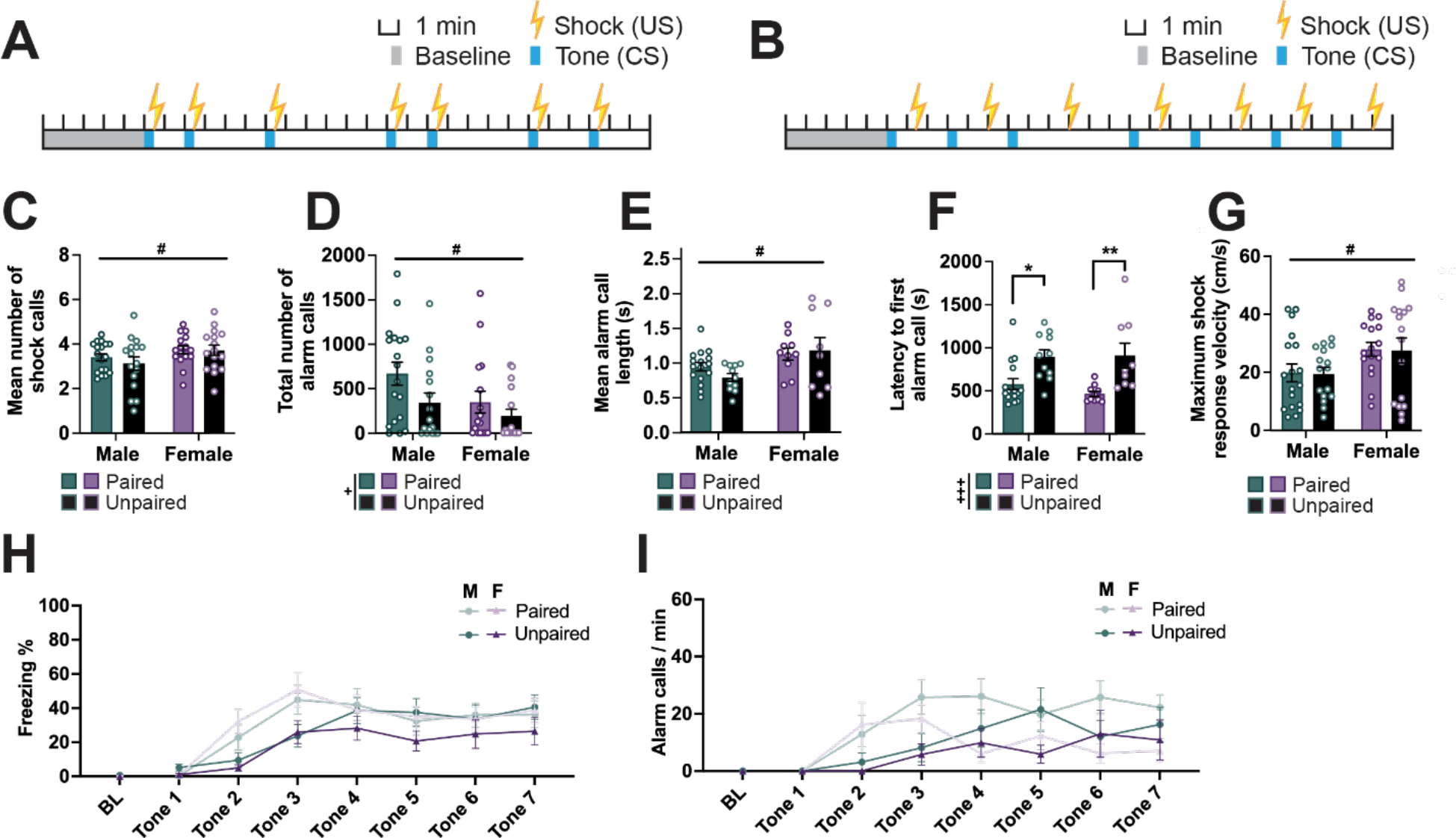
Unpaired CS and US results in delayed latency to alarm call. **A-B**. Schematics of cued fear conditioning showing timing and duration of tones (CS) and shocks (US) for paired (**A**) and unpaired (**B**) protocols. **C**. Bar graph showing the mean number of shock calls emitted in response to each shock averaged across the trial (7 shocks) by each animal. **D**. Bar graph depicting the total number of alarm calls emitted during a fear conditioning trial. **E**. Bar graph depicting the mean alarm call length. **F**. Bar graph depicting the latency of each animal to emit their first alarm call. **G**. Bar graph depicting the maximum velocity reached in response to each shock averaged across all 7 shocks within a trial by each animal. **H**. Line graph showing the percent of time male and female rats within each protocol group (paired vs unpaired) spent freezing during baseline (BL, first 2 minutes) and each tone. **I**. Line graph showing the normalized (per minute) alarm call rate of male and female rats within each protocol group (paired vs unpaired) during baseline (BL, 5 minutes) and each tone. Ns: C, D, G - I: Paired = 18 males, 16 females, Unpaired = 16 males, 16 females; E & F: Paired = 16 males, 10 females, Unpaired = 11 males, 9 females (Non-alarm callers excluded). Bar graphs depict mean ± SEM, and each dot represents a single animal. Symbols along line graphs indicate mean ± SEM. Significant main effects of pairing (+) and sex (#), and post hoc comparisons (*) are denoted with different symbols, with either 1, (*p* < 0.05), 2 (*p* < 0.01) or 3 (*p* < 0.001) symbols depicting degree of significance.

Stimulus delivery was controlled and videos of the animals’ behavior recorded using Ethovision (v16, Noldus Technologies) and infrared digital cameras mounted on top of each conditioning chamber. Time spent freezing, the occurrence of darts, and the maximum velocity reached as response to the shock were analyzed using freely available Python-based software ScaredyRat (Mitchell et al., 2022). An animal was classified as a Darter if they performed one or more darts (movement at a speed exceeding 20 cm/s) during one or more tones (excluding the first two tones as well as the immediate response to the shock). For all fear conditioning trials, baseline freezing was recorded from the first 2 minutes of the 5-minute stimulus-free time in the chamber. Freezing was defined in ScaredyRat as the absence of observable movement, with a minimum bout duration of 1 s.

Our data set includes animals (20 males, 20 females) that were part of a different project involving systemic injections of clozapine-N-oxide (CNO) for circuit-specific activation using designer receptors activated by designer drugs (DREADDs). These rats underwent intracranial infusions of viral vectors (AAV_retro_ pmSyn1-EBFP-Cre, and either pAAV_8_-hSyn-DIO-hM4D(Gi)-mCherry, pAAV_8_-hSyn-DIO-hM3D(Gq)-mCherry, or pAAV_8_-hSyn-DIO-mCherry, all sourced from Addgene) under isoflurane anesthesia, allowing 5-6 weeks of recovery prior to behavioral testing. However, most (35/40) of these animals did not show any fluorescent signal marking viral expression. To ensure that this experience did not influence behavioral outcomes, we compared these animals to same-sex animals receiving the same foot shock intensity (0.5 mA) but no surgical experience or CNO exposure. This analysis revealed no differences in alarm call rate, length or latency, or in shock call rate (Extended Figure 2-1), thus justifying their inclusion in the data set. These animals were also exposed to extinction learning and retrieval (Fig 6). Extinction learning was carried out 24 hours after fear conditioning, and extinction retrieval 24 hours after extinction learning. For both tests, animals were brought into the testing room 30 min before the start of the test to habituate. The testing room and chambers were the same as for fear conditioning, but with different lighting (8 lux), scent (2-4 drops of Dr Bronner’s peppermint-scented pure-castile liquid soap placed on a train underneath the test cage floor) and chamber features (black plexiglass floor covering the metal grids). For extinction learning, baseline behavior was recorded for the full 5 minutes of stimulus-free time prior to the start of tone presentations. For these experiments we included the full baseline duration because we observed that animals that emitted alarm calls prior to tone start frequently did so also at the end of the baseline period. Thus, to be able to compare freezing and alarm-calling, we chose to record both for the full duration. Animals received a total of 20 presentations of the same tone CS they heard during fear conditioning, at varying intervals (90-330 seconds), with no shocks. Similarly, for extinction retrieval 24 hours after extinction learning the animals were exposed to a baseline period of 5 min followed by 3 presentations of the same tone at varying intervals (150-240 seconds). Time spent freezing in extinction learning and retrieval was quantified using a combination of ScaredyRat and hand-scoring by trained investigators (each tone evaluated by two independent investigators and their scores averaged, and where the scores differed by more than 2 seconds a third investigator resolved the discrepancy).

### USV recording and analysis

Throughout the behavioral experiments we recorded vocalizations emitted in the audible and ultrasonic range (0-120 kHz, sampling rate 250 kHz) using microphones (CM16, Avisoft Bioacoustics, Germany) mounted over each conditioning chamber. Pilot testing showed that each microphone was only able to detect sounds from the chamber it was in, no cross-detection from other chambers was observed (data not shown). The audio files were processed with DeepSqueak (v3, Coffey et al., 2019), a publicly available user interface which uses machine learning to detect spectrograms typical of rodent USVs. All detected calls were manually confirmed by a trained investigator. Temporal alignment of the audio and behavioral data was confirmed by observation of the tone cue within the audible range of the spectrogram. We then aligned each detected call with an epoch (Baseline, Tone, Shock or ITI) and identified alarm calls based on specific criteria (call length ≥ 70 ms, main frequency of the call ≤ 30 kHz, and change in frequency ≤ 10 kHz).

Group differences were analyzed using SPSS (v27) and GraphPad Prism (v8). The specific test used was determined by data type and structure, see Results for details. Geisser-Greenhouse correction for non-sphericity was applied when needed. Correlation analyses (Fig 6 and 7) were conducted in SPSS using Pearson’s method if both variables in the analysis were normally distributed (determined using the Shapiro-Wilk test), and Spearman’s if one or both were non-normally distributed. Bonferroni correction was used to adjust *p*-values for multiple comparisons where appropriate, and an alpha level of 0.05 was used throughout. Outlier data points were excluded based on the ROUT method in Graphpad Prism (FDR *q* = 0.1).

## Results

### Sex differences in short ultrasonic and audible calls during fear conditioning as a function of shock intensity

To investigate the nature of USVs occurring during cued fear conditioning (Fig 1A), we recorded and analyzed auditory data ranging from 0 to 120 kHz. During the baseline period, while the rats explore the conditioning apparatus prior to any tones or shocks, animals frequently engage in various types of chatter in the form of short high-frequency calls (Fig 1B). Females emitted more such calls than males (independent t-test, *t* = 2.355, *p* = 0.0200, Fig 1D). Another specific type of call was observed when the animals were given the foot shocks, and these will be referred to as “shock calls” (Fig 1C). To assess how well the shock calls reflect the potential degree of discomfort the animals experience, we asked whether shock intensity influences their occurrence. A two-way ANOVA suggested a significant main effect of shock intensity (F_2, 128_ = 14.36, *p* < 0.001), with both males and females receiving 1 mA foot shocks emitting the highest number of these calls (Males 0.3 mA vs 1 mA: *p* < 0.001; Males 0.5 mA vs 1 mA: *p* < 0.001; Females 0.3 mA vs 1 mA: *p* = 0.040, Fig 1E).

In line with past research, we observe robust freezing behavior across the fear conditioning trial in both males and females (Fig 1F-G). At the end of the trial (during Tone 7) females froze significantly less than males (two-way ANOVA, main effect of Sex: F_1, 128_ = 3.995, *p* = 0.048, Fig 1H). Additionally, the main effect of shock intensity on freezing during Tone 7 was significant (F_2, 128_ = 4.034, *p* = 0.020), with a significant post hoc contrast only in Male 0.3 mA vs 0.5 mA group comparison (*p* = 0.006).

### Sex differences in alarm calls during fear conditioning as a function of shock intensity

A distinct type of USV is emitted when rats experience aversive events, such as maternal separation as pups (Granata et al., 2022; Wöhr & Schwarting, 2008), and exposure to fear conditioning (Borta et al., 2006; Wöhr et al., 2005; Yee et al., 2012) or a context associated with a predator odor (Fendt et al., 2018) in adults. These calls are relatively long with a principal frequency of 22 kHz (Fig 2A), and will be referred to here as “alarm calls”. Animals exposed only to handling, the experimental apparatus and tones without foot shocks do not emit alarm calls, suggesting novelty and the stress of performing the experiment alone are not sufficient to elicit them (Extended Data 2-1). Typically these calls were not emitted until after the second tone-shock pairing, although males receiving 0.5 mA or 1 mA foot shocks started emitting alarm calls earlier than those receiving 0.3 mA shocks (two-way ANOVA, main effect of Shock intensity: F_2, 81_ = 8.092, *p* < 0.001; post hoc comparisons Male 0.3 mA vs 0.5 mA *p* = 0.004, Male 0.3 mA vs 1 mA *p* < 0.001, Fig 2B). Overall males made significantly more alarm calls than females (two-way ANOVA, main effect of Sex: F_1, 128_ = 16.67, *p* < 0.001, main effect of Shock intensity: F_2, 128_ = 7.622, *p* < 0.001), and males receiving strong foot shocks made more alarm calls than those receiving mild shocks (Fig 2C). Alarm calls occurred consistently also during the inter-trial intervals (ITIs), and they were of similar length between epochs across the whole trial (Extended Data 2-1). Mean alarm call length did not differ between sexes, but there was a significant interaction between sex and shock intensity (two-way ANOVA, F_2, 81_ = 3.226, *p* = 0.045) driven by females in the 0.5 mA condition making significantly shorter alarm calls than those in the 0.3 mA condition (*p* = 0.018 Fig 2D). The pattern of alarm calling across the fear conditioning session mirrors that of freezing (Fig 1F-H), and at the last tone there was a significant main effect of shock intensity with stronger shocks associating with higher alarm call rates (two-way ANOVA, F_1, 128_ = 19.53, *p* < 0.001) and a trend towards a main effect by sex (males emitting more alarm calls than females, F_2, 128_ = 2.912, *p* = 0.058, Fig 2E-G).

### Unpredictability of the shock drives delay in initiation of alarm calls

Next, we asked if predictability of the US affects the nature of USVs by comparing male and female rats exposed to paired (co-terminating US and CS, Fig 3A) and unpaired (independently occurring US and CS, Fig 3B) fear conditioning (shock intensity = 0.5 mA for all). Rats exposed to unpaired fear conditioning did not differ from those exposed to paired fear conditioning on measures of the number of shock calls (Fig 3C), alarm call length (Fig 3E) or maximum shock response velocity (Fig 3G) as assessed by two-way ANOVA. However, on each of these measures we observed a main effect of sex, with female rats making more shock calls (F_1, 62_ = 4.092, *p* = 0.047), and longer alarm calls (F_1, 62_ = 5.235, *p* = 0.027), and moving faster after the shock (F_1, 62_ = 6.825, *p* = 0.011). We found a main effect of pairing condition on both total number of alarm calls emitted and the latency to alarm; rats in the unpaired condition emitted overall fewer alarm calls (main effect of Pairing: F_1, 62_ = 4.608, *p* = 0.036; main effect of Sex: F_1, 62_ = 4.625, *p* = 0.035, Fig 3D), possibly due to their longer first alarm call latencies (F_1, 62_ = 20.500, *p* < 0.001, Fig 3F and I). The effect on latency is unlikely to be explained by the variations in timing of shock delivery between pairing conditions (Fig 3A-B). Both males and females in the paired condition on average start emitting alarm calls after the second shock occurring at 7.5 min (average latency 9.6 min for males, 7.6 min for females), while in the unpaired condition both start emitting alarm calls after the third shock occurring at 14 min (average latency 14.9 minutes for both sexes). In other words, on average one additional shock exposure was required to elicit alarm calls in the unpaired versus paired condition. Interestingly, there is also a delay in reaching peak freezing in the unpaired condition compared to the paired one (Fig 3H).

### Darting does not associate with differences in USVs

While freezing is to date the most commonly quantified index of learning the CS-US association, it is by no means the only available and informative motor behavior rodents engage in. Darting refers to rapid movements occurring during the CS, particularly after a number of CS-US pairings have been established (Greiner et al., 2019; Gruene et al., 2015; Hersman et al., 2020; Mitchell et al., 2022). We classified all animals as either Darters or Non-darters based on the occurrence of one or more darts (movement exceeding a speed of 20 cm/s) during tones 3-7 (Colom-Lapetina et al., 2019; Gruene et al., 2015; Mitchell et al., 2022), and observed Darters across all shock intensities (Males: *X*^2^ = 0.224, *p* = 0.894; Females: *X*^2^ = 1.048, *p* = 0.592, Fig 4A). We observed more Darters among females than males (*X*^2^ = 5.877, *p* = 0.015). Two-way ANOVAs were conducted to investigate whether Darters differed from Non-darters within sex on USV variables (baseline chatter, shock calls, alarm calls, alarm call length, and alarm call latency, Fig 4B-F), and no significant main effects or interactions were observed (*p*s > 0.1701). By contrast, other motor behaviors (maximum velocity reached in response to the shock, freezing) differed significantly as a function of darting. Both male and female Darters had faster maximum shock response velocities (main effect of Darting: F_1, 130_ = 15.93, *p* < 0.001; Male Darters vs Non-darters *p* = 0.043; Female Darters vs Non-darters *p* < 0.001), in addition to a main effect of sex (F_1, 130_ = 14.21, *p* < 0.001, Fig 4G). There was a significant main effect of darting on freezing across the fear conditioning trial (F_1, 130_ = 9.428, *p* = 0.003), with female Darters freezing significantly less than female Non-darters (*p* = 0.043) and a trend in the same direction for males (*p* = 0.065, Fig 4H).

**Figure 4.**
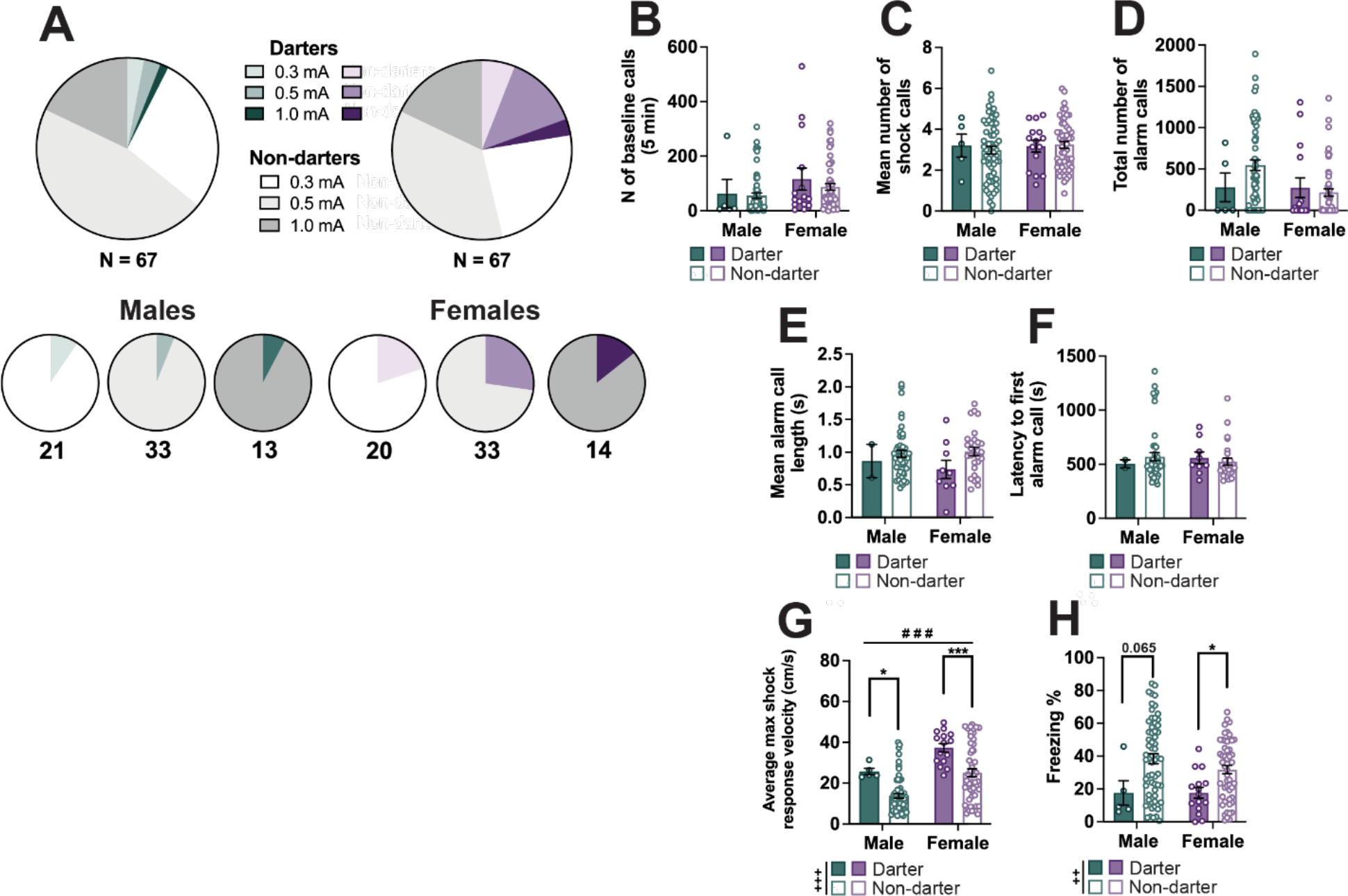
Darting does not associate with differences in USV features in either sex. **A**. Pie charts showing the proportion of Darters and Non-darters across the whole male and female cohorts (top row) and separately for each shock intensity group (bottom row). Numbers underneath each chart denote the number of animals included within the chart. **B**. Bar graph showing the total number of baseline chatter calls emitted by male and female Darters and Non-darters prior to fear conditioning. **C**. Bar graph showing the mean number of shock calls emitted in response to each shock averaged across the trial (7 shocks) by each animal. **D**. Bar graph depicting the total number of alarm calls emitted during a fear conditioning trial. Two-way ANOVA suggests no significant main effects or interactions. **E**. Bar graph depicting the mean alarm call length. **F**. Bar graph depicting the latency of each animal to emit their first alarm call. **G**. Bar graph depicting the maximum velocity reached in response to each shock averaged across all 7 shocks within a trial by each animal. **H**. Bar graphs showing the percentage of time animals spent freezing across all 7 tones of a trial. Ns: B-H: Males = 5 Darters, 62 Non-darters; Females = 15 Darters, 52 Non-darters. Bar graphs depict mean ± SEM, and each dot represents a single animal. Significant main effects of darting (+) and sex (#), and post hoc comparisons (*) are denoted with different symbols, with either 1, (*p* < 0.05), 2 (*p* < 0.01) or 3 (*p* < 0.001) symbols depicting degree of significance.

### Alarm calling distinguishes high and low freezing only in males

We noted that in addition to inter-individual variability in alarm call rate, some rats did not emit any alarm calls during the whole fear conditioning trial. To explore whether this behavior constituted a behavioral phenotype, we compared alarm call emitting rats (Alarm callers) to rats that did not emit a single alarm call (Non-alarm callers) on other USV and motor behaviors. A chi-squared test indicated that shock intensity groups differed in the frequency of Alarm callers (Fig 5A), with a significant effect in females (*X^2^* = 6.758, *p* = 0.034) and a trend in males (*X^2^* = 5.838, *p* = 0.054). In both sexes we observed more Alarm callers in high shock intensity groups (1 mA) than the lower shock intensity groups. We also replicated the alarm call analyses depicted in Fig 2 after excluding rats that did not emit any alarm calls, and the key findings remained unchanged (Extended Figure 5-1). While the data shown in Figure 4 is collapsed across shock intensity and analyzed by two-way ANOVA (Sex x Alarm calling), we also performed linear regression analysis to evaluate the contribution of shock intensity to group differences (Sex, Shock intensity and Alarm calling as predictors). First we found a significant Sex x Alarm calling interaction effect on the number of baseline calls (F_1, 130_ = 8.933, *p* = 0.003, Fig 5B), with a significant pairwise comparison only in females (*p* = 0.002) suggesting that Alarm callers are more vocal during the baseline period. In the linear regression model, Sex was the strongest and the only significant predictor of baseline calling (β 0.230, *p* = 0.009), with a trend towards a significant contribution of Alarm calling (β = −0.149, *p* = 0.100). Next, a two-way ANOVA suggests main effects of both Sex and Alarm calling on the number of shock calls (main effect of Sex: F_1, 130_ = 6.517, *p* = 0.012; main effect of Alarm calling: F_1, 130_ = 12.820, *p* < 0.001, interaction: F_1, 130_ = 7.891, *p* = 0.006, Fig 5C), with a significant pairwise comparison only in males where Alarm callers emitted significantly more shock calls (*p* < 0.001). The linear regression analysis suggests that a larger portion of this effect was attributable to Shock intensity (β = 0.371, *p* < 0.001) than Alarm calling (β = −0.166, *p* = 0.047), although both served as significant predictors. No significant differences were observed between Alarm callers and Non-alarm callers on shock response velocity (Fig 5D). Interestingly, when analyzing the average percentage of the CS duration the rats spent freezing, we found a significant main effect of Alarm calling (F_1, 130_ = 17.91, *p* < 0.001; Sex x Alarm calling interaction: F_1, 130_ = 16.32, *p* < 0.001) and a significant pairwise comparison within males (*p* < 0.001), suggesting that rats that did not make any alarm calls also froze considerably less than their conspecifics that did emit alarm calls. Linear regression suggests that this effect could not be attributed to Shock intensity (β = 0.106, *p* = 0.213), but was significantly affected by Alarm calling (β = −0.266, *p* = 0.003).

**Figure 5.**
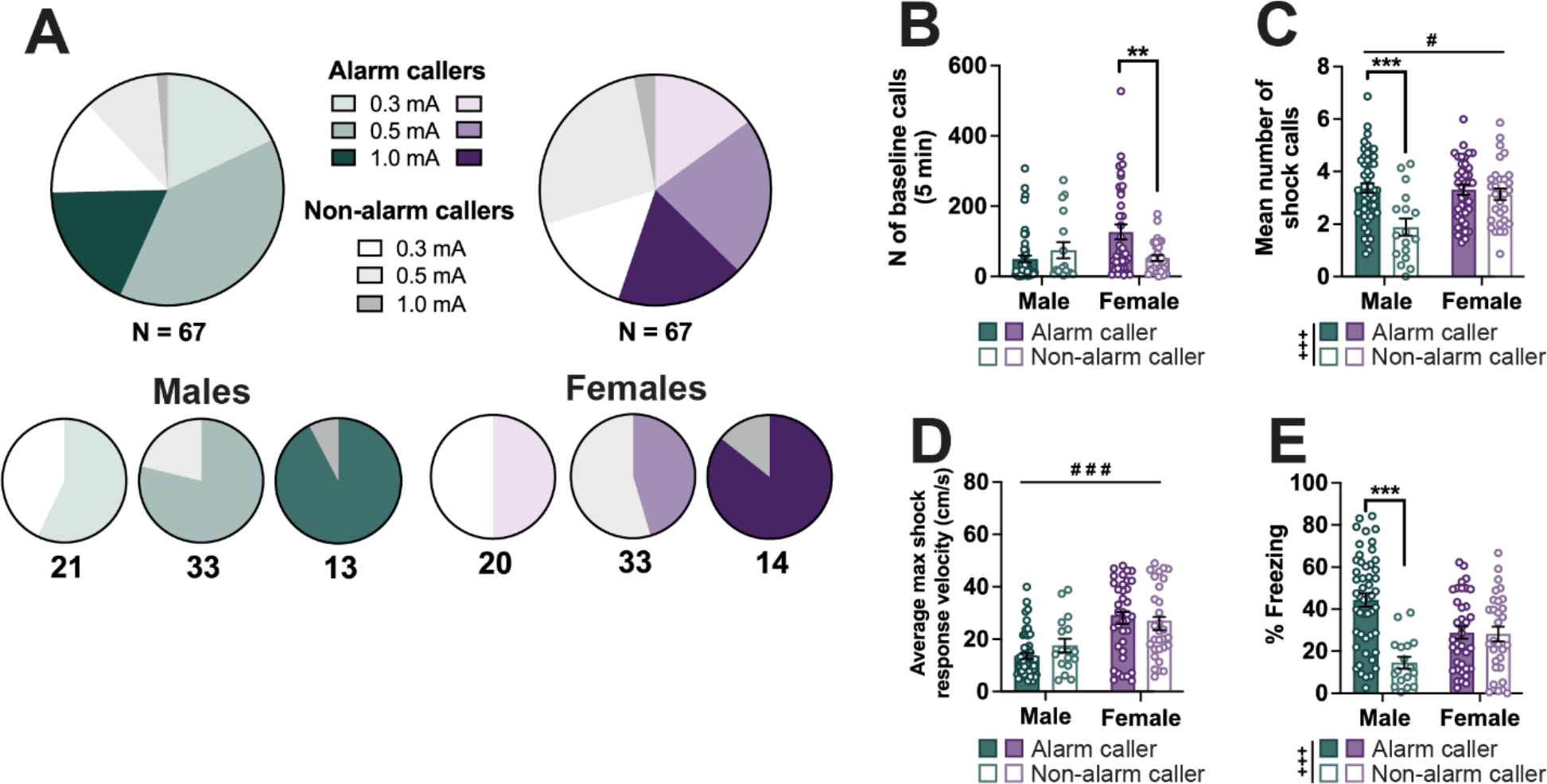
Tendency to emit alarm calls as a dichotomous phenotype associates with freezing in males only. **A**. Pie charts showing the proportion of Alarm callers and Non-alarm callers across the whole male and female cohorts (top row) and separately for each shock intensity group (bottom row). Numbers underneath each chart denote the number of animals included within the chart. **B**. Bar graph showing the total number of baseline chatter calls emitted by male and female Alarm callers and Non-alarm callers prior to fear conditioning. **C**. Bar graph showing the mean number of shock calls emitted in response to each shock averaged across the trial (7 shocks) by each animal. **D**. Bar graph depicting the maximum velocity reached in response to each shock averaged across all 7 shocks within a trial by each animal. **E**. Bar graphs showing the percentage of time animals spent freezing across all 7 tones of a trial. Ns: B - E: Males = 50 Alarm callers, 17 Non-alarm callers; Females = 37 Alarm callers, 30 Non-alarm callers. Bar graphs depict mean ± SEM, and each dot represents a single animal. Significant main effects of alarm caller status (+) and sex (#), and post hoc comparisons (*) are denoted with different symbols, with either 1, (*p* < 0.05), 2 (*p* < 0.01) or 3 (*p* < 0.001) symbols depicting degree of significance.

### Alarm calls are extinguished across sexes, but correlate with freezing more strongly in males

To explore how dynamically USV emissions change as the animals acquire and extinguish the CS-US association, we recorded USVs and behavior throughout fear conditioning (FC), extinction learning (EL) and extinction retrieval (ER). Using a two-way repeated measures ANOVA we found that while the amount of baseline calls did not change across these trials (Fig 6A), their nature changed in that some animals were observed emitting alarm calls during the baseline period (Fig 6B) leading up to EL (2/20 males, 3/20 females) and ER (4/20 males, 0/20 females). Mirroring this, there was a significant main effect of trial on the latency to first alarm (mixed effects model, effect of Trial: F_1.878, 29.11_ = 18.64, *p* < 0.001; Trial by Sex interaction: F_2, 31_ = 5.040, *p* = 0.013, Fig 6C). Post hoc contrasts were only significant in males (FC vs EL *p* < 0.001; FC vs ER *p* < 0.001), with a trend observed in females (FC vs EL *p* = 0.055). We also observed that alarm calls emitted across these trials got progressively longer (mixed effects model, main effect of Trial: F_1.581, 25.30_ = 7.799, *p* = 0.004, Fig 6D) with significant post hoc comparisons only in males (FC vs EL *p* = 0.032; FC vs ER *p* = 0.026). The overall rate of alarm calling reduced over time (two-way repeated measures ANOVA, main effect of Trial: F_1.555, 59.09_ = 12.50, *p* < 0.001; main effect of Sex: F_1, 38_ = 9.534, *p* = 0.004, Fig 6E) with significant post hoc comparisons in males (FC vs EL *p* = 0.004; FC vs ER *p* = 0.003). Looking at the pattern of alarm calls across each trial (Fig 6F’, F’’ and F’’’), female alarm calls consistently peaked at a similar time point to males, but remained at a lower level throughout each trial (two-way repeated-measures ANOVA, main effects of Sex in FC [F_1, 38_ = 13.05, *p* = 0.001], EL [F_1, 38_ = 7.350, *p* = 0.01] and ER [F_1, 38_ = 4.875, *p* = 0.033]). During FC we also observed higher levels of freezing in males than females (two-way repeated-measures ANOVA, main effect of Sex: F_1, 38_ = 6.572, *p* = 0.014). In ER, by contrast, females froze more than males (two-way repeated-measures ANOVA, main effect of Sex: F_1, 38_ = 7.702, *p* = 0.009), with no sex differences observed in EL. While male alarm call emission sloped downward during EL, it was still observed during the last 5 tones in 5 out of 20 of rats (compared to only 1 female rat). The following day during ER male alarm call emission was also considerably more prevalent (12/20 males, 3/20 females), and remained so for each of the three tone presentations. Across all trials, we found significant or trending correlations between alarm call rate and freezing in males (Fig 6H), but not in females (*p*s > 0.658, Fig 6I).

Next, we asked whether the patterns of motor and vocal behaviors in one trial correlated with motor or vocal behaviors in consequent trials. Full correlation heatmaps are presented in Figure 7. The heatmaps are split into quadrants as follows: top quadrants show motor behaviors (freezing, shock response velocity) correlated with later motor (left) or vocal behaviors (right), while bottom quadrants show vocal behaviors correlated with later motor (left) and vocal behaviors (right). In both sexes, but more notably in males, we see consistency between alarm calls across FC, EL and ER in the form of significant correlations between alarm call rates and latencies (bottom-right quadrants, Fig 7A-F). In males, we see alarm and shock call parameters in FC correlating with freezing during early EL (Tone 1, Fig 7A bottom-left), while in females early EL freezing is predicted by FC freezing (Fig 7D, top-left). Additionally, in males we observe EL freezing to be positively correlated with ER alarm call rate (Fig 7C, top-right).

**Figure 6.**
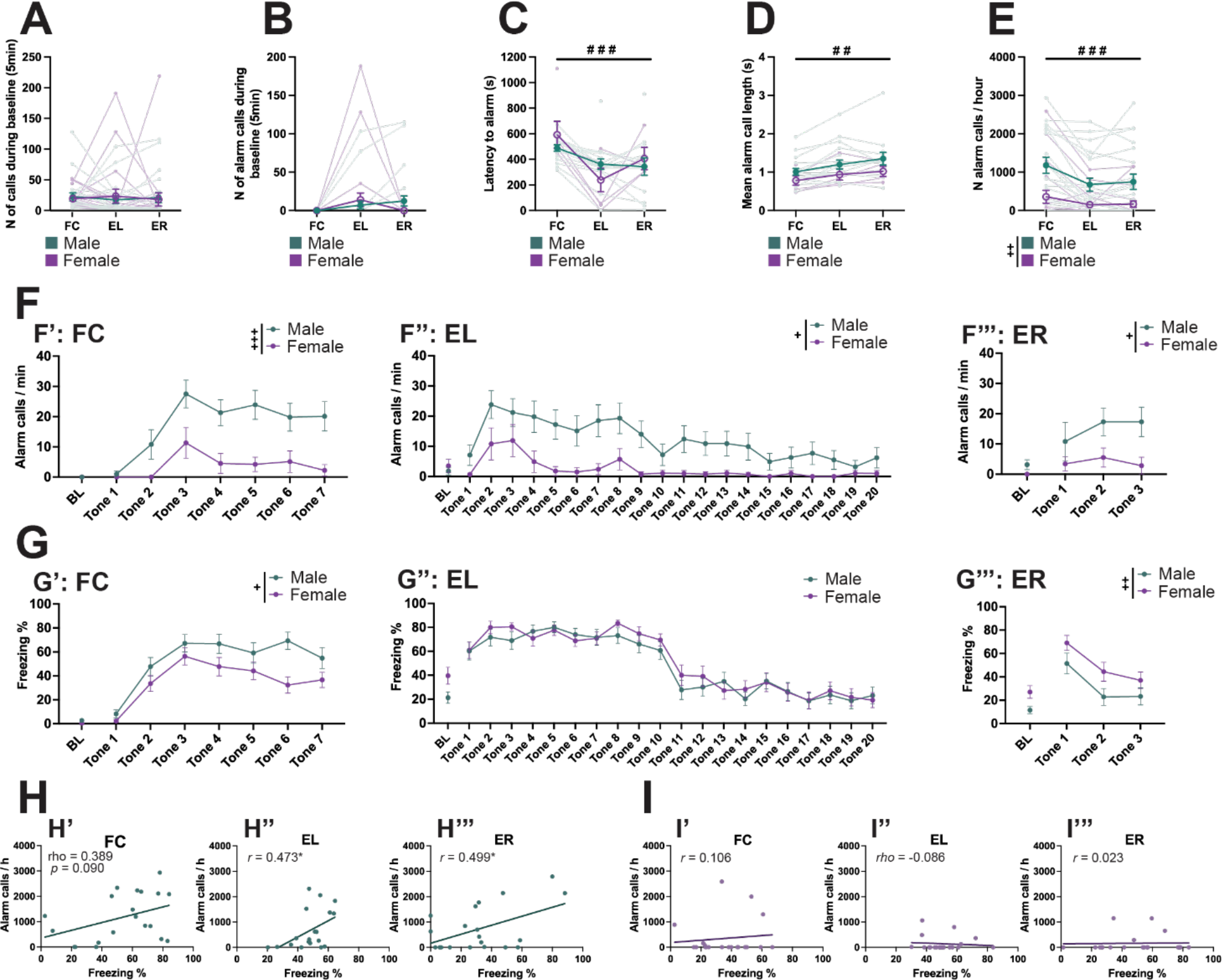
Sex differences in extinction of alarm calling. **A-E**. Connected scatter plots of the number of all USVs emitted during baseline (**A**), the number of alarm calls emitted during baseline (**B**), the latency to first alarm (**C**), the mean alarm call length (**D**), and the rate of alarm calls normalized to 1 hour (**E**) of male and female rats during fear conditioning (FC), extinction learning (EL) and extinction retrieval (ER). Light colors represent individual animals, while dark colors with error bars represent sex means. **F**. The rate of alarm calls, normalized per 1 minute, during the baseline (BL) and each tone presentation during FC (**F’**), EL (**F’’**) and ER (**F’’’**), shown separately for males and females. **G**. Percent of time spent freezing during the first 2 minutes of BL for FC (**G’**), and the first 5 minutes of BL for EL (**G’’**) and ER (**G’’’**). **H-I**. Scatter plots and regression lines showing the within-trial correlation of freezing (x-axis) and alarm call rate (y-axis) for each trial (‘ = FC, ’’ = EL, ’’’ = ER), shown separately for males (**H**) and females (**I**). Correlation coefficients (*r* = Pearson’s r, *rho* = Spearman’s rho) are shown inside each panel. Ns: A, B, E - I: 20 males, 20 females; C & D: 20 males, 8 females (excluding animals who made no alarm calls in any trial). Dark-toned data points depict mean ± SEM. Significant main effects of sex (+) and trial type (#) are denoted with different symbols, with either 1, (*p* < 0.05), 2 (*p* < 0.01) or 3 (*p* < 0.001) symbols depicting degree of significance.

**Figure 7.**
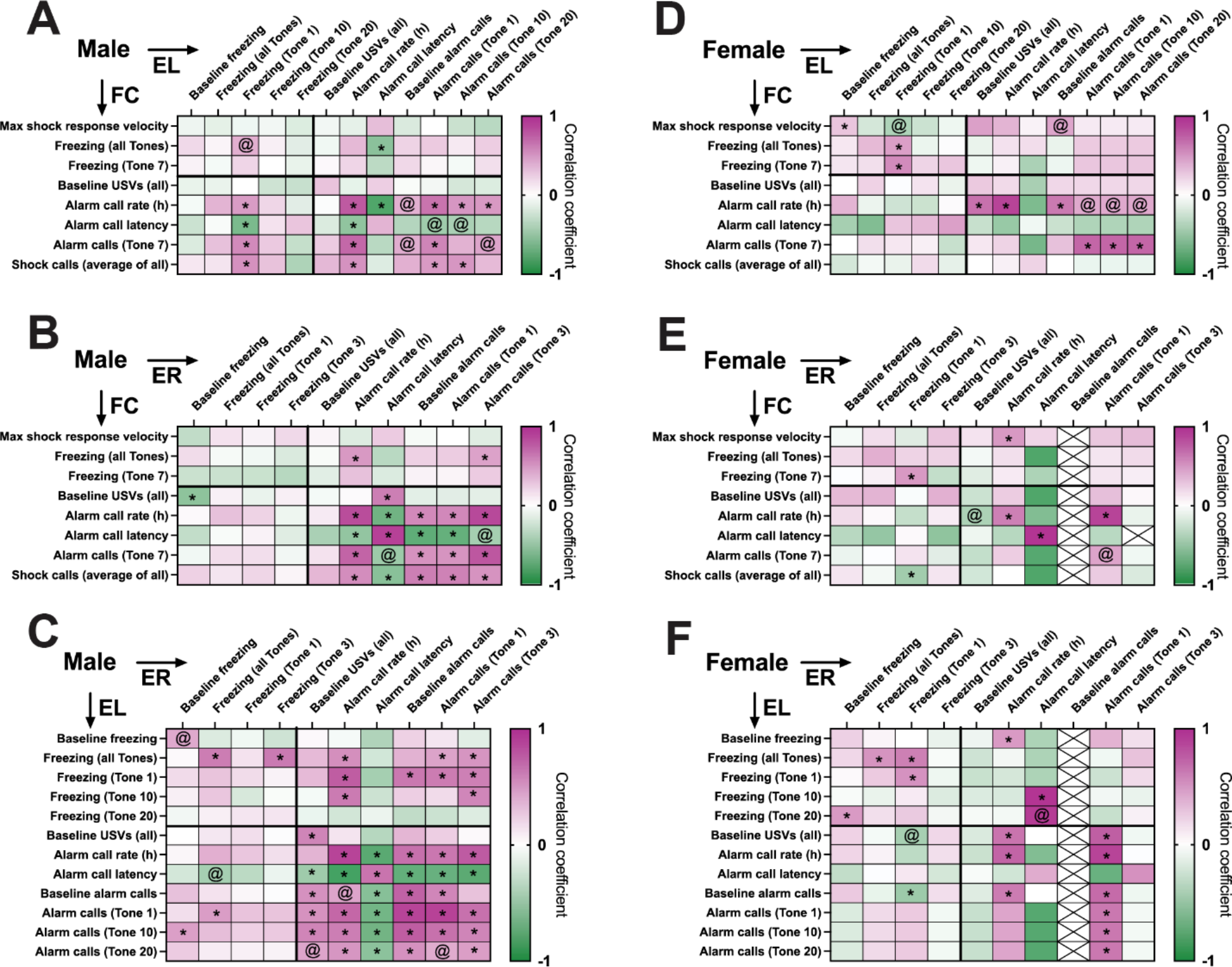
Correlation patterns of freezing and alarm call rates across fear conditioning (FC), extinction learning (EL) and retrieval (ER). **A-F**. Heatmaps showing correlation coefficients (Pearson’s or Spearman’s depending on the distribution of the data in each variable pair) of motor and vocal behaviors during one of the trials (y-axis) and one of the subsequent trials (x-axis), separately for males (**A-C**) and females (**D-E**). Color scale corresponds to correlation coefficient and direction (magenta = positive, green = negative). Significant (uncorrected *p* < 0.05) correlations are marked by *, trending (uncorrected *p* < 0.1) correlations are marked by @.

## Discussion

Across these studies we provide a rich picture of rat USV production during fear conditioning, demonstrating how it relates to other behavioral readouts of internal states, and sex differences therein. Alarm calls in male rats were more frequent than in female rats, tracked more robustly with the intensity of the aversive experience (shock intensity), and were more resistant to extinction. Additionally, male rats that abstained from alarm vocalizations also had markedly low levels of freezing during fear conditioning. Across testing sessions we observed significant correlations in alarm call rates in both males and females, suggesting that it may represent a trait-like individual characteristic. In males we also saw alarm call rates and freezing correlating within and across sessions. Darting did not associate with any of the measured USV parameters in either sex. In both males and females the experience of foot shocks that were not predictably preceded by a tone resulted in a delay in alarm call initiation compared to those exposed to tone-paired foot shocks. We also found that in both sexes short, audible calls occurred immediately after the foot shock, scaling with shock intensity. Multidimensional behavioral measurement such as this – particularly strategies that include sex and other individual differences – will push our field towards more valid and successful translational research.

We found that during the baseline period female rats emitted more short, high-frequency calls than males did. These calls have been extensively reported on in appetitive circumstances (Knutson et al., 2002; Reinhold et al., 2019; Schwarting et al., 2007; Takahashi et al., 2010; Wright et al., 2010), drug-exposure (Knutson et al., 1999; Lawson et al., 2021; Simola, 2015; Wright et al., 2010) and preclinical models of autism spectrum disorder (Caruso et al., 2020), and their production is modified by drug-withdrawal (Lin et al., 2018), sleep-deprivation and lithium (Wendler et al., 2019). The majority of these experiments use exclusively male subjects, although some suggest female mice are less likely to spontaneously emit such vocalizations (Michael et al., 2020). Our work highlights the need to include females in such experiments, at least when using a species or strain in which females reliably emit these calls, to fully understand how high-frequency calls relate to affective states and respond to interventions. It is plausible that they reflect variable states or nuances thereof across sex, and without including females the conclusions drawn will only capture part of the picture.

Audible calls, akin to those recorded in mice during a tail suspension test (Ruat et al., 2022), were observed in response to the foot shock in our paradigm. Such calls have been reported in female rats experiencing fear conditioning (Schwarting, 2018). Interestingly, the number of shock calls emitted scaled with the intensity of the foot shock in both males and females. Unlike alarm calls, we also observed that all rats (except for one male rat) emitted at least some shock calls. While we do not have access to the animals’ subjective perception of pain or discomfort on exposure to the shock, our findings suggest that vocalizations in direct response to painful stimuli could serve as a proxy for intensity, as opposed to alarm calls which emerge with a delay and show sex-biased scaling with intensity. Further work should explore whether similar dose-response curves can be observed with other modalities of aversive stimuli, to determine if this response is selective to the electric foot shocks used here. Another interesting question is: are these calls modulated by pharmacological interventions targeting the pain system, such as opioid-antagonist naloxone (as shown before for alarm calls in males, Oliveira & Barros, 2006)? Our work shows the value in recording these commonly unreported vocalizations as a potential window into the rodents’ experience of aversion and pain.

Overall, we observed that male rats made more alarm calls than females across all experiments. In males, shock intensity had a stronger effect on the total number of alarm calls emitted, as well as the latency to initiate alarm calls, than in females. Categorically, males were more likely than females to emit alarm calls, and those males that did so also froze significantly more during fear conditioning. Taken together, these findings point to alarm calls likely serving as a more accurate metric of negative affective state, such as perceiving a threat or experiencing discomfort, in male rats than in females. Other reports have also been published suggesting males emit more alarm calls than females (Willadsen et al., 2021), and in a study involving only males the authors found a similar dose-response relationship with shock intensity as we observe here (Wöhr et al., 2005). Similar to our findings, these groups also reported a lack of alarm calling in some animals, along with a correlation between freezing and alarm call durations (Willadsen et al., 2021; Wöhr et al., 2005). Here we show this correlation with freezing extends to the number of alarm calls emitted, but only in males. Our findings imply that the propensity to emit alarm calls may be a part of a broader threat response phenotype in males, although a potential genetic basis for the Alarm caller phenotype has not yet been investigated. As the rat stock used for these experiments is outbred, the contribution of genetic variability to our findings is plausible. In an illustration of this possibility, knock-out of the serotonin transporter gene reduces alarm call rates in male and female Wistar-crossed rats (Willadsen et al., 2021). It would also be intriguing to investigate whether Alarm callers and Non-alarm callers differ from each other in some other features, such as neuroanatomy or neuronal ensembles engaged during fear conditioning. Studying such divergent response styles could help elucidate individual differences in factors that affect response to traumatic events, and recovery thereof, with translational relevance for human psychiatric disorders.

We also observed alarm calls in several, although not all, rats throughout extinction learning and retrieval. The occurrence of these calls during an extinction or test session following fear conditioning has been reported before (Kikusui et al., 2001; Wöhr et al., 2005) in male Wistar rats, and here we expand on this by demonstrating the pattern of alarm calls across training sessions and sexes. Similar to freezing, we see alarm calls at a low level during the baseline recording time of extinction learning, with a sharp increase during tone CS presentation, followed by a gradual decay. This decay in alarm call rate was considerably steeper in females, while no differences were observed in terms of freezing during this trial. This finding highlights an important caveat in behavioral tests: our conclusions depend critically on our choice of readout. If alarm call rate was conceptualized as a measure of associative learning (discussed in more depth below), this data set would point to a remarkable sex difference in extinction learning and retrieval, with females outperforming males on both. But if in the same animal cohort we only had access to freezing data we would conclude that no sex differences in EL could be observed, and if anything the females show worse extinction retrieval than males. As it stands, our findings cannot be used to determine which behavior is more suitable for measuring internal states such as fear. Rather, we call for caution in operationalizing any internal state as a singular behavioral output. Rapid technological advances in behavioral recording and *in vivo* interrogation of neuronal activity help us take large strides towards new discoveries about what makes individuals acquire and extinguish aversive memories. However, much also remains to be done in terms of deep understanding of what the behaviors we routinely record mean, and how to best harness them for translational aims.

As certain behaviors during fear conditioning, such as darting, are known to predict later performance on extinction retrieval (Gruene et al., 2015), an important question going forward is whether alarm calling could serve as a similar predictor. In males much more strongly than females, we observed correlations between alarm call rates and freezing both within a trial and across fear conditioning and extinction. For example, we observed that alarm call rate during fear conditioning was positively correlated with freezing in extinction learning, but only during the start of it. This could suggest that in males alarm call rate relates to the strength of association between the CS and US. However, alarm calls during fear conditioning were not related to freezing at the end of extinction learning or across extinction retrieval, suggesting this behavior may not predict the success of extinction. We do also see alarm calls in some rats during the baseline period of extinction trials, suggesting either generalization or stress-sensitization, once again more so in males than females. Additionally, in males freezing during EL correlated with alarm call rate during ER. These findings align with the idea of behavior as a circular-causality loop, as opposed to an arc with linear and replicable outputs occurring after certain inputs (Gomez-Marin & Ghazanfar, 2019). What the animal experiences and perceives affects its behavior, and that behavior further affects what it perceives and how it behaves, all within the context of individual history and characteristics. This framework of the dynamic nature of behavior expression fits with our findings; rather than universal associations between outputs (behaviors) and inputs (experience of shocks), we observe variable relationships which are further moderated by individual factors like sex and a trait-like tendency to emit alarm calls.

An important question to consider in case of any behavior occurring in the context of aversive learning is to what extent the behavior is associative. One of the key utilities of Pavlovian conditioning is the acquisition, and later extinction, of a learned association between the US and the CS. Currently the expression of such learning is primarily gauged by observing behavior, such as freezing and darting. However, it should be noted that no behavior specifically and exclusively denotes associative CS-US learning. Freezing, considered the gold standard of measuring associative learning, occurs in response to not just the CS but the context in which learning has taken place (Kamprath & Wotjak, 2004), and in most standard fear conditioning protocols it is challenging to distinguish the proportion of freezing driven by associative and nonassociative components. Freezing has also been shown in response to a CS unpaired from the US (Cossio et al., 2016; Hersman et al., 2020; Trott et al., 2022) although some report observing very little to no freezing specifically during the CS in such conditions (Maren, 2000; Maren et al., 2001). Our findings suggests that alarm calls, at least to the depth measured here, may reflect an associative learning component of fear conditioning, but by no means do so exclusively. We do see a similar pattern between alarm call emission and freezing, i.e. a gradual rise in alarm call rate as the animals experience more CS-US pairings, and a recurrence followed by gradual decay during extinction. The fact that emission of alarm calls rarely starts after just one foot shock suggests that it is not merely a response to acute discomfort, and thus could be influenced by learning in addition to continuity of the discomfort. Furthermore, when the association between the US and CS was reduced by carrying out unpaired fear conditioning, there was a delay in alarm call initiation, also suggesting that predictability or a learned association with a predictor may have played a role in alarm call emission. However, we also robustly observe alarm calls during unpaired fear conditioning and during ITIs regardless of CS-US pairing, indicating that they are not tied specifically to the tone, and thus not uniquely indexing the associative component of learning. Alarm call recording does not have as long a history as freezing as a measure of threat learning, and many crucial control experiments remain to be conducted such as explicit comparison between contextual and cued conditioning, sensitivity of alarm calls to habituation, and tests of long-term recall. Our data and that of others is foundational for building an understanding of how to best make use of recorded vocalization in studies of aversive memories.

While important from an ethological perspective, alarm calls may not signal the same experiences in male and female rodents. Alarm call emission in male rats is largely in line with prior literature, ergo it tracks with stimulus intensity and defensive motor behaviors. However, in female rats alarm calls were observed largely independent of stimulus intensity and defensive behaviors. Further research is needed to understand which factors within female rodents affect the nuances in USV production in order to best utilize this behavior as a readout in behavioral experiments. Significant inter-individual variability (from none at all to thousands of calls within a trial) as well as intra-individual stability (as evidenced by correlation across trials) argue for studies employing USVs to favor a within-subjects design, as opposed to cross-sectional approaches. Investigating the source of these individual differences, such as what makes an Alarm caller, could also be fruitful for understanding different threat or stress response types. Our findings show that USVs are a valuable, non-invasive source of data that is sensitive to experimental manipulations, but what they tell us about the animals’ affective states may depend on several variables, including sex.

## Acknowledgments

We thank Lauren Granata (Heather Brenhouse lab, Northeastern University) for help setting up the ultrasonic vocalization recording system.

