## Supplementary Figures for "Sounding the alarm: sex differences in rat ultrasonic vocalizations during Pavlovian fear conditioning and extinction"

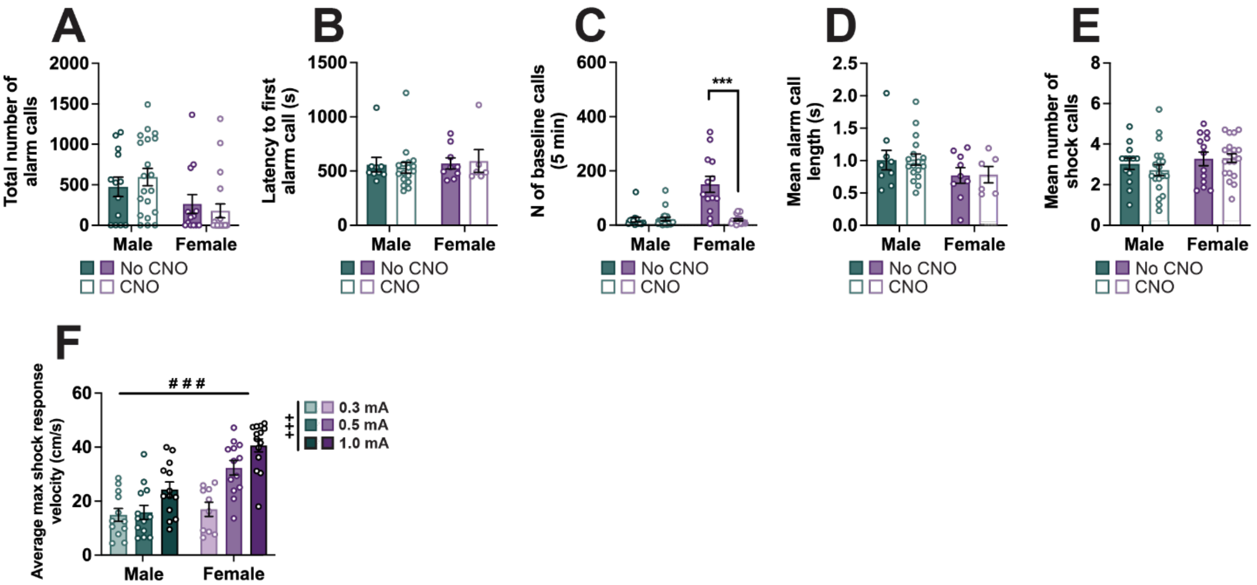


**Extended Data 1-1. CNO exposure does not affect alarm call parameters in males or females 1. A-E.** Bar graphs depicting the total number of alarm calls emitted during fear conditioning (**A**), the latency to the first alarm call during the session (**B**), the number of all calls emitted during the baseline period (**C**), mean alarm call length (**D**) and the average number of shock calls emitted per animal across all 7 shocks (**E**) of male and female rats exposed to 0.5 mA foot shocks, compared between experiments (one involving clozapine-N-oxide [CNO] injection and one with no injections). **F.** Bar graphs depicting the average peak velocity reached immediately after shock delivery, split by sex and shock intensity.

Bar graphs depict mean ± SEM, and each dot represents a single animal. Significant main effects of shock intensity (+) and sex (#), and post hoc comparisons (*) are denoted with different symbols, with either 1, ($p < 0.05$), 2 ($p < 0.01$) or 3 ($p < 0.001$) symbols depicting degree of significance.

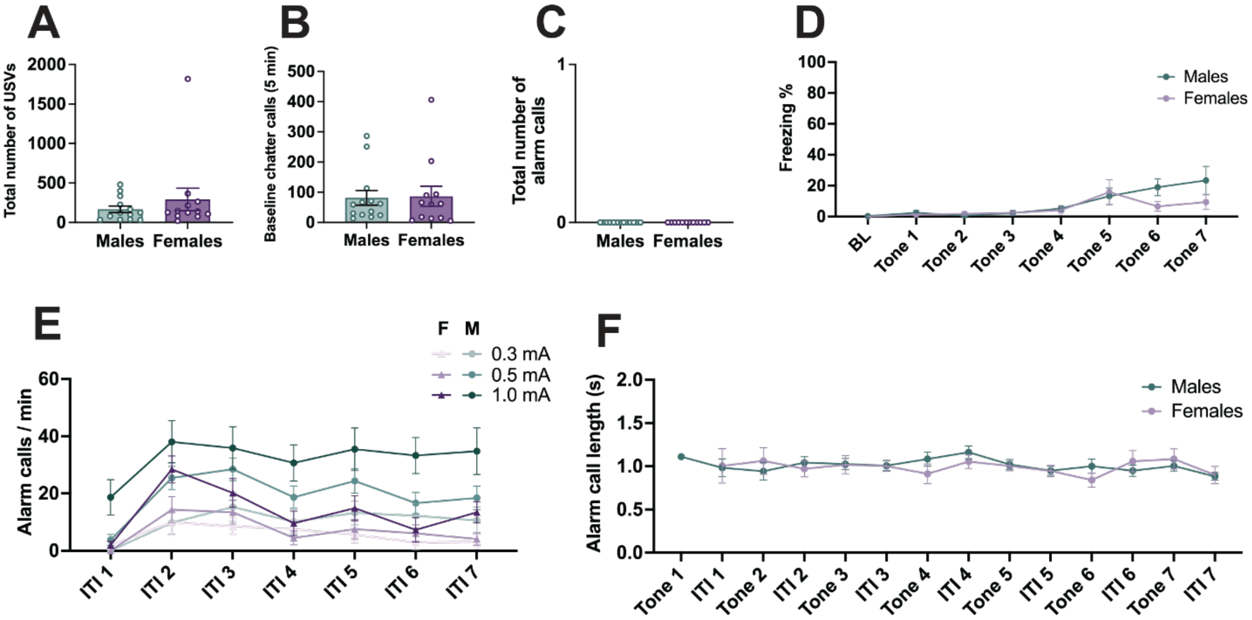


**Extended Data 2-1. Mere exposure to the experimental context and tone are not sufficient to produce alarm calls, and shock-driven alarm calls are also observed during the inter-trial intervals (ITIs). A-C.** Bar graph showing the total number of USVs across the whole trial (**A**), total number of USVs emitted during the baseline (**B**), and total number of alarm calls (**C**) emitted by male and female rats exposed to the same testing procedure, chamber and tones as those in Figure 1, but with no foot shocks. **D.** Line graph showing the percent of time spent freezing during baseline (first 2 minutes only) and each tone of animals exposed to only the tones without shocks. **E.** Line graph depicting the rate of alarm calling of male and female rats in each shock intensity group (0.3 mA, 0.5 mA and 1mA) during the inter-trial intervals (ITIs). **F.** Line graph showing the mean alarm call length as measured during each tone and ITI, separately for males and females. Ns: A-D: 13 males, 12 females; E: 67 males (0.3 mA = 21, 0.5 mA = 33, 1 mA = 13), 67 females (0.3 mA = 20, 0.5 mA = 33, 1 mA = 14), F: 50 males, 37 females (Non-alarm callers excluded).

Bar graphs depict mean ± SEM, and each dot represents a single animal. Symbols along line graphs indicate mean ± SEM.

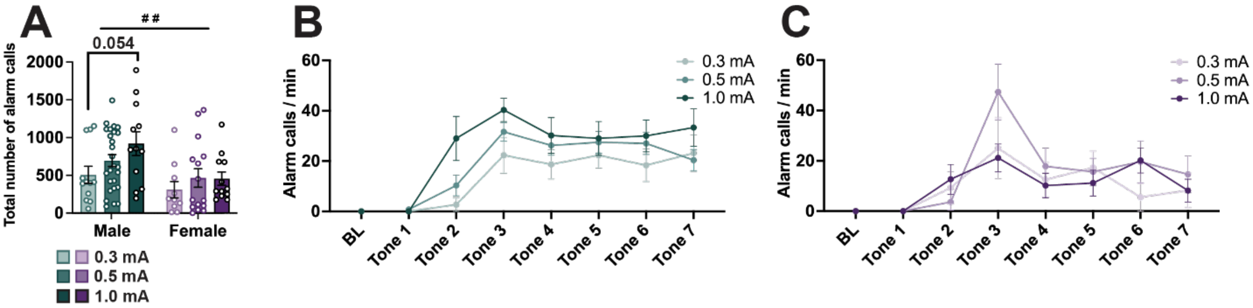


**Extended Data 5-1. Removing Non-alarm callers does not eliminate the sex difference in alarm call rate. A.** Total number of alarm calls emitted by rats included in Fig 2C, excluding rats that did not make any alarm calls. **B-C.** Line graphs showing the normalized (per minute) alarm call rate of male (**B**) and female (**C**) rats across shock intensity groups during baseline (BL, 5 minutes) and each tone, excluding rats that did not make any alarm calls. Ns: 50 males (0.3 mA = 12, 0.5 mA = 26, 1 mA = 12), 37 females (0.3 mA = 10, 0.5 mA = 15, 1 mA = 12).

Bar graphs depict mean ± SEM, and each dot represents a single animal. Symbols along line graphs indicate mean ± SEM. Significant main effects of shock intensity (+) and sex (#), and post hoc comparisons (*) are denoted with different symbols, with either 1, ($p < 0.05$), 2 ($p < 0.01$) or 3 ($p < 0.001$) symbols depicting degree of significance.
